# Distinct activation mechanisms of β-arrestin 1 revealed by ^19^F NMR spectroscopy

**DOI:** 10.1101/2023.03.06.531433

**Authors:** Ruibo Zhai, Zhuoqi Wang, Zhaofei Chai, Conggang Li, Changwen Jin, Yunfei Hu

## Abstract

β-Arrestins (βarrs) are functionally versatile proteins that play critical roles in the G-protein-coupled receptor (GPCR) signaling pathways. While the classical theory of GPCR-mediated βarr activation centers around the formation of a stable complex between βarr and the phosphorylated receptor tail, emerging evidences highlight the indispensable contribution from membrane lipids for many receptors. Due to the intrinsic complexity of βarr conformational dynamics, detailed molecular mechanisms of its activation by different binding partners remain elusive. Herein we present a comprehensive study of the structural changes of βarr1 in critical structural regions during activation using ^19^F NMR method. We demonstrate that phosphopeptides derived from different classes of GPCRs show distinct abilities in inducing βarr1 activation. We further show that the membrane phosphoinositide PIP_2_ independently modulates βarr1 conformational dynamics without displacing its autoinhibitory carboxyl tail, leading to a distinct partially activated state. Our results delineate two activation mechanisms of βarr1 by different binding partners, uncovering a highly multifaceted conformational energy landscape for this protein family.

## INTRODUCTION

The ubiquitously expressed β-arrestins (βarrs) play essential roles in G-protein-coupled receptor (GPCR) signaling, involving not only receptor desensitization ^1,2^ but also receptor internalization and the initiation of diverse G protein-independent signaling cascades ^3,4^. The presence of only two βarrs (βarr1and βarr2, also known as arrestin-2 and arrestin-3) in the human genome and the fact that they respond to over 800 GPCRs suggest high structural adaptability. The increasingly recognized functional versatilities and the intrinsic structural plasticity of βarrs have urged for a deeper understanding of their activation mechanisms.

A commonly believed model of βarr activation by GPCRs depicts a biphasic binding process with the formation of a ‘tail-engaged’ complex in which βarr binds only the receptor phosphorylated tail, and a ‘core-engaged’ complex in which βarr further associates with the receptor transmembrane core ^5,6^. However, this scenario is challenged by several lines of evidences that have begun to surface. Firstly, phosphorylation of receptor tail is not always required for βarr binding ^7,8^, and the “catalytic” activation of βarrs reveals a receptor tail-independent process that relies on membrane phosphatidylinositol 4,5-bisphosphate (PIP_2_) ^9^. Secondly, the recent work by Janetzko et al ^10^ demonstrates the categorizing of GPCRs into two groups, one requiring PIP_2_ for βarr recruitment and the other not, which strongly coincides with the historical classification of the A and B classes depending on whether they show transient (class A) or tight (class B) interactions with βarrs ^4,11^, Thirdly, among the GPCR-βarr1 complex structures available thus far, all receptors harbor a class B tail that is either native (e.g. the native V2 vasopressin receptor, V2R ^12^ and neurotensin receptor 1, NTSR1 ^13,14^) or chimeric (e.g. β1-adrenoceptor, β1AR ^15^ and M2 muscarinic receptor, M2R^16^, both fused with the phosphorylated V2R tail, a model peptide and the strongest binder to βarrs ^5^), whereas no stable complexes were obtained for native class A tails. Therefore, the molecular basis underlying the difference between class A and B receptors, as well as the mechanism of PIP_2_-mediated βarr activation, remain to be understood.

We herein report a ^19^F NMR study of βarr1 conformational dynamics during activation by receptor-derived phosphopeptides and PIP_2_. We show that while the phosphopeptide from V2R (V2Rpp) sufficiently displaces the autoinhibitory carboxyl tail of βarr1 and induces dramatic conformational changes in various regions, the phosphopeptides from the class A β2-adrenergic receptor (β2AR) are ineffective in promoting βarr1 activation. We further demonstrate that PIP_2_ independently triggers partially activation of βarr1 via a distinct mechanism without displacing its carboxyl tail. Our results provide a dynamic overview of βarr1 conformational changes during activation at residue level, and highlight a complex conformational landscape of βarr1 that can be differentially modulated by multiple binding partners.

## RESULTS

### Design of βarr1 constructs for site-specific ^19^F NMR studies

To obtain site-specific structural information by ^19^F NMR, we started from a fulllength functional cysteine-less βarr1 construct ^17^ (referred to as βarr1 hereafter) and introduced individual cysteine mutations at desired sites for chemical ligation of ^19^F probe (**Supplemental data Fig. S1**). A total of 20 sites were selected covering essential regions in both the N- and C-domains (**Fig. 1a**). All mutants maintain structural and functional integrity as verified by ^1^H NMR and clathrin binding assay.

**Figure 1.**
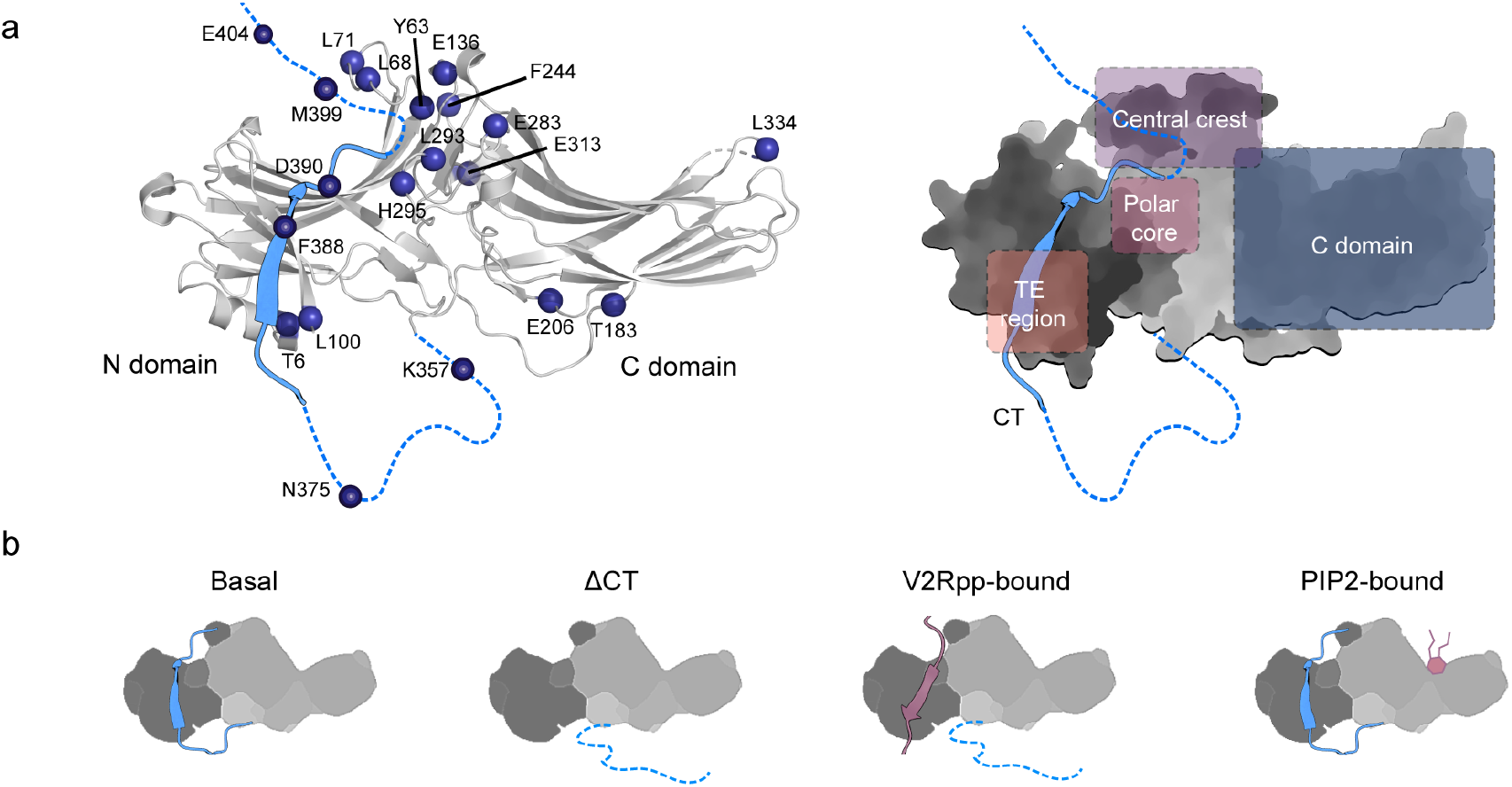
^19^F labeling of βarr1 for NMR experiments. (**a**) Selected residues for ^19^F labeling in the inactive βarr1 crystal structure (PDB: 1JSY, left panel) and schematic illustration of the key structural regions associated with βarr1 activation (right panel). The middle region of CT is shown as ribbon colored in blue, whereas the proximal and distal regions unobservable in the crystal structure are shown as blue dashed lines. (**b**) Illustration of the four functional states of βarr1 examined in this study.

On one hand, to probe the βarr1 conformational changes induced by receptor phosphorylated tails, particularly the classical V2Rpp, we focused on structural regions undergoing critical changes during activation. These include the carboxyl tail (hereafter abbreviated as CT or βarr1-CT to distinguish from the receptor C-terminal tail), the three-element (TE) interacting site, the polar core and the central crest region based on available structures ^18–21^. On the other hand, for investigation of changes induced by PIP_2_, which binds in the C-domain, a number of additional sites in this domain were chosen.

In this work, we examine the conformational dynamics of βarr1 in four functional states, namely the autoinhibited basal state represented by the full-length βarr1, the CT-released pre-activated state represented by the βarr1-ΔCT construct (residues 1-382), and the V2Rpp- or PIP_2_-bound states prepared using the full-length βarr1 protein (**Fig. 1b**). In the following sections, we will analyze the differential effects of receptor tails and PIP_2_ binding on βarr1 conformational dynamics.

### V2Rpp-induced conformational changes of βarr1

Since V2Rpp activates βarr1 by displacing its autoinhibitory CT, we compared the ^19^F NMR data of βarr1 in the basal, ΔCT and V2Rpp-bound states to trace its conformational changes during activation. The data suggest that while all states show a certain degree of structural dynamics, the V2Rpp-bound state exhibit the highest heterogeneity with many sites displaying two or more conformations (**Supplemental data Fig. S2a**). In particular, residues in the central crest adopt a uniform conformation in the basal state, while they show multiple conformations when activated. Moreover, the chemical shift differences (Δδ) can reflect the amplitude of conformational (or environmental) changes, and we observe prominent changes for residues in the polar core, central crest and the CT regions, indicating significant alterations of the local structures (**Supplemental data Fig. S2b**). Detailed changes are analyzed below.

#### The CT and TE regions

The βarr1 CT comprises over 60 residues (K354-R418) and can be divided into the proximal, middle and distal regions. Both the proximal and distal regions are invisible in the crystal structures, and the middle region forms the ‘s20’β-strand according to the gpcrdb.com numbering. Among the six sites selected, F388C and D390C are in the middle region, whereas K357C, N375C and M399C, E404C are in the proximal and distal regions, respectively (**Fig. 1a**). Upon V2Rpp binding, we observe significant chemical shift perturbations accompanied by linewidth reduction for both middle and distal regions, but not in the proximal region (**Figure 2a-b**). Particularly, the F388C and D390C sites show the largest linewidths (98 and 87 Hz) in the basal state, reflecting their tight binding to the N-domain and thus slower tumbling rate, whereas the values decrease to 41 and 52 Hz in the V2Rpp-bound state, reflecting the release of the CT. The linewidths of the M399C and E404C sites are smaller than the middle region residues in the basal state, consistent with their more distal locations and thus higher 5 flexibility. Upon V2Rpp-binding, both sites show a major peak with an upfield chemical shift change and decreased linewidth, and a minor peak with chemical shift similar to the basal state. Therefore, even when the middle region of the CT is completely displaced by V2Rpp, a small population of the distal region may still transiently interact with the N-domain.

**Figure 2.**
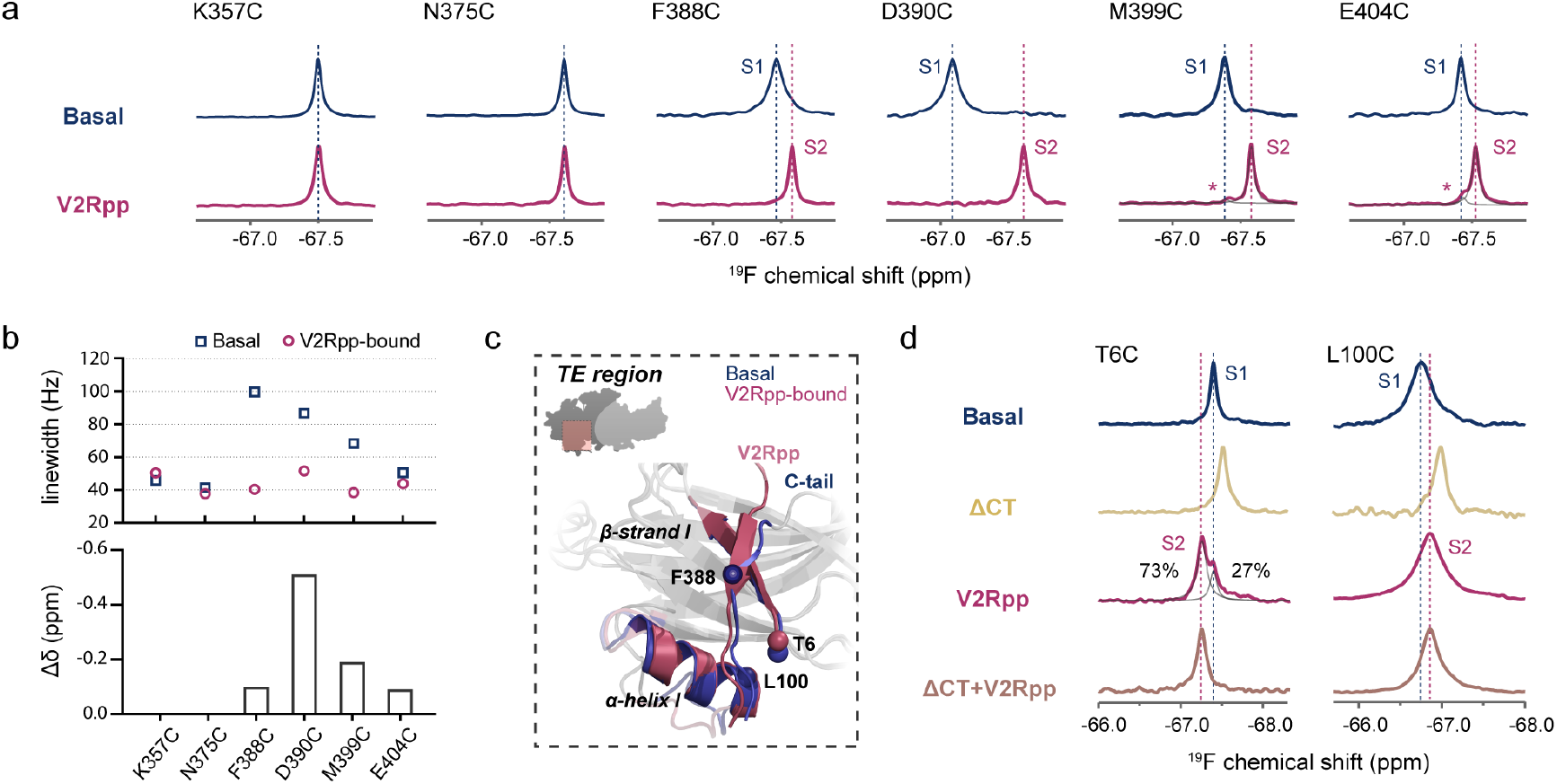
^19^F NMR spectra of βarr1 CT and TE region in different functional states. (**a**) ^19^F NMR spectra of residues in the CT region in the basal and V2Rpp-bound states. Asterisk indicates the minor peak observed in the distal region. (**b**) Linewidths and chemical shift differences of the CT region in the basal and V2Rpp-bound states. (**c**) Local structure of the TE region showing the location of the ^19^F labeling sites. The inactive (PDB: 1JSY) and V2Rpp-activated (PDB: 4JQI) states crystal structures of βarr1 are shown in blue and magenta, respectively. (**d**) ^19^F NMR spectra of the T6C and L100C sites in the TE region in different functional states.

The TE interaction region is one essential structural “lock” that holds βarr1 in the inactive state, at which the CT packs with the β-strand I and α-helix I in the N-domain. Apart from the F388C and D390C sites in the CT described above, we also used two labeling sites, T6C and L100C, in the N-domain to monitor conformational dynamics in this region (**Fig. 2c**). Similar trends of chemical shift changes are observed for both sites. CT removal leads to upfield chemical shift changes, consistent with increased shielding effect when negative charges in the CT are absent, whereas V2Rpp-binding shifts the resonances back towards downfield positions, consistent with the decreased shielding induced by the phosphate groups (**Fig. 2d**). However, differences in local dynamics are also observed for the two sites. For one thing, the L100C site shows extremely broad resonance linewidths (~ 247 and 257 Hz in the basal and V2Rpp-bound states), reflecting intermediate exchange among multiple conformations on the NMR timescale (μs-ms) and implying intrinsic heterogeneity at the tip of α-helix I. For another, the T6C sites shows two peaks (S1 and S2) in the V2Rpp-bound state with populations of ~ 27% and 73%, whereas only a single S2 peak is observed for the V2Rpp-bound βarr1-ΔCT sample. This indicates that the S1 state, which has the same chemical shift with the basal conformation, may actually represent a subpopulation in which the CT is not fully displaced. Note that the V2Rpp molar concentrations used in these experiments are 1.2-fold that of βarr1, and is sufficient to induce close to 100% CT displacement as observed at both the F388C and D390C sites. Because T6 locates at the bottom side of the TE region and is closer to the N-terminal end of the s20 strand, a possible explanation is that although the central part of the s20 strand is fully displaced by V2Rpp (**Supplemental data Fig. S3**), its N-terminal region can remain partially in contact with the N-domain.

#### The polar core

The polar core, formed by a network of electrostatic interactions between the gate loop and surrounding residues, is another essential structural element that locks the N- and C-domains into the inactive orientation. Activation is accompanied by a flip of the gate loop and disruption of the polar core, allowing inter-domain twisting. Two sites in the gate loop, L293C and H295C, were chosen to monitor the local conformational changes (**Fig. 3a**). Different from other regions, both sites show conformational heterogeneity in the basal state, displaying two peaks with populations of ~80-90% (S1) and 10-20% (S2) (**Fig. 3b**). However, we observe different changes of dynamics for the two sites.

**Figure 3.**
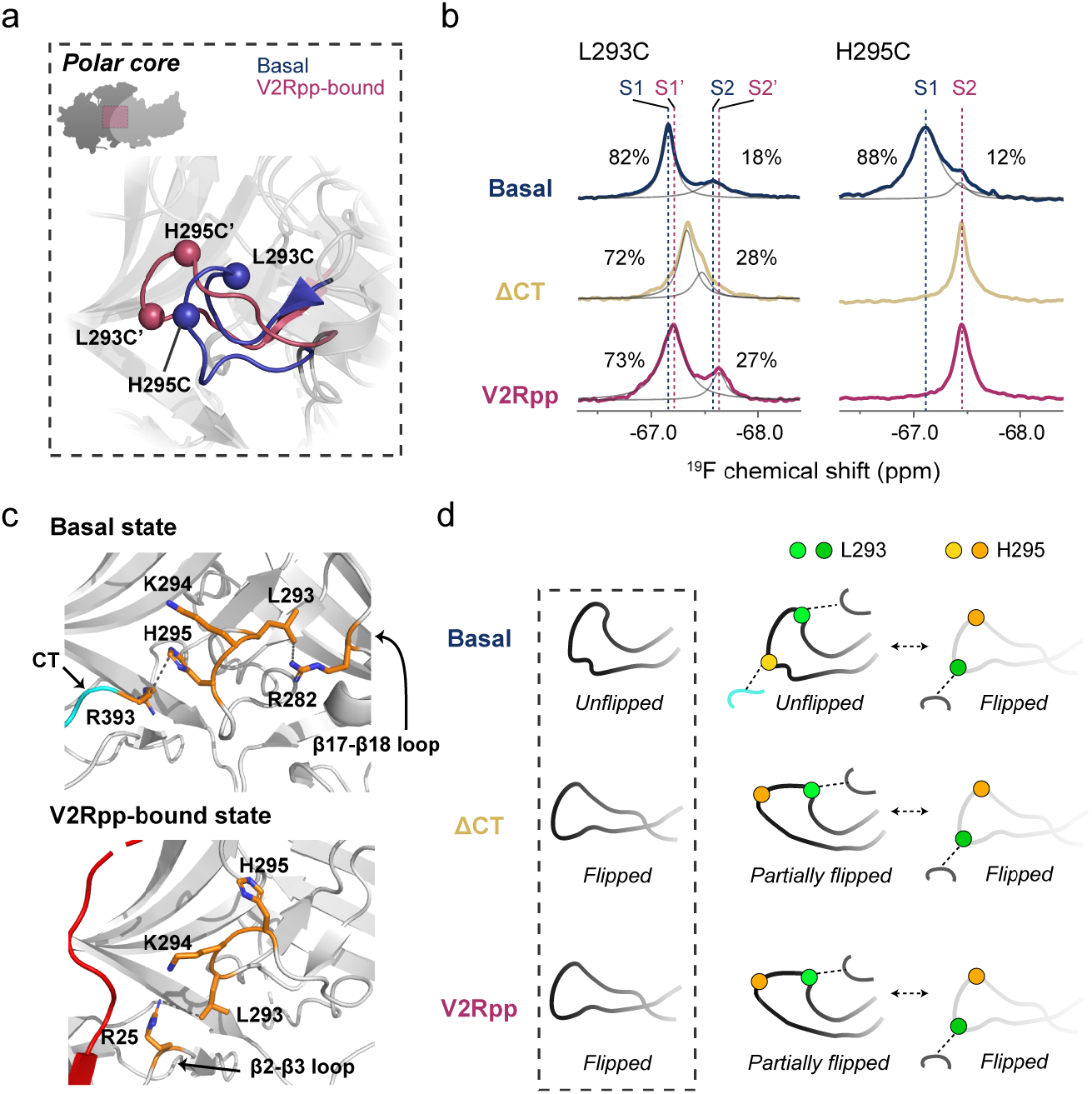
Conformational changes of the gate loop probed by ^19^F NMR. (**a**) ^19^F labeling sites in the gate loop mapped onto the inactive (PDB: 1JSY, blue) and V2Rpp-activated (PDB: 4JQI, magenta) βarr1 crystal structures. (**b**) ^19^F NMR spectra of the gate loop residues in the three functional states. (**c**) Detailed local conformations of the gate loop in the basal (PDB: 1G4M) and V2Rpp-bound (PDB: 4JQI) states. The observed close contacts between H295-R393, L293-R282 in the basal state, and K294-R25 in the V2Rpp-bound states are shown in dashed lines. (**d**) A schematic illustration showing the complex changes of gate loop conformations during activation inferred from the ^19^F NMR data. The autoinhibitory CT is colored in cyan. For comparison, the dashed box shows the static “unflipped” and “flipped” conformations observed in crystal structures ^18^–^20^. The “flipped” conformation in the ΔCT functional state is deduced from the crystal structure of the pre-activated visual arrestin splice variant p44 ^21^.

In the case of H295C, the linewidth of the peak S1 (230 Hz) is notably larger than that of L293C (113 Hz) in the basal state, suggesting that the H295C site exhibit more significant conformational exchanges on the intermediate NMR timescale (μs-ms). Both CT removal and V2Rpp binding result in a single peak for H295C at the S2 position with decreased linewidth (~ 115 Hz and 127 Hz), indicating that the S2 peak represents the active “flipped” conformation as observed in the crystal structures ^18–21^. Thus, the S1 peak most probably corresponds to the inactive conformation in which H295 shows close contact with R393 from the CT, and release of the CT results in a full transition into the active state in which the H295 side chain becomes relatively flexible (**Fig. 3c & Supplemental data Fig. S4**).

The L293C site, however, shows more complex spectral changes. Dual conformations are observed for this site in all three states, suggesting this site can intrinsically adopt two alternative conformations. CT release results in the decrease of chemical shift difference between the two peaks, which may be attributed to a faster exchange rate or a more similar chemical environment between the two conformations. Binding to V2Rpp leads to the separation of the two peaks (S1’ and S2’) again, but with slightly altered chemical shift values compared to the basal state, and a similar population distribution as in the ΔCT state. Inspection of the crystal structures shows two distinct conformations for L293. Its side chain points rightward to contact the β17-β18 loop in the basal state, while it points leftward to contact the β2-β3 loop in the V2Rpp-bound state (**Fig. 3c & Supplemental data Fig. S4**). We speculate that the S1 and S2 (or the S1’ and S2’) peaks correspond to the β17-β18 loop-contacting and β2-β3 loop-contacting conformations respectively, and that βarr1 activation induces an increased preference for the latter conformation. The small chemical shift differences between the S1-S1’ and S2-S2’ peak pairs may reflect subtle variations in local environments for both conformations in the two functional states.

Based on the different spectral behaviors of the two sites, the gate loop probably occupies a more complex conformational energy landscape than the simple two-state “flipping” suggested by the crystal structures. We suggest that the L293C spectra may more closely reflect the existence of dual conformations in the gate loop in all functional states, whereas the complete absence of the S1 peak for H295C in both ΔCT and V2Rpp-bound states may be due to the lack of the contacting R393 residue and the resulting flexibility of H295C once the CT is displaced. Thus, we propose a modified mechanism of the gate loop conformational changes during activation as shown in **Fig. 3d**. It exists as an equilibrium between the “unflipped” inactive conformation and the “flipped”active conformation in the basal state, whereas CT displacement releases the H295 residue and results in a “partially flipped” intermediate conformation, in which L293 remains contacting the β17-β18 loop. The “partially flipped” conformation interchanges with the “flipped” active conformation in both the ΔCT and V2Rpp-bound states. Although we cannot rule out the possibility that spin labeling may affect local dynamics, spectral differences between the three functional states clearly reveal that βarr1 activation promotes the “flipped” active conformation, demonstrated by the increased populations of the S2 peaks for both L293C and H295C labeling sites.

#### The central crest

The central crest, in particular the finger loop, is the key interaction site with the receptor core. V2Rpp binding disrupts a number of charge interactions and enable the finger loop to flip up into its active form. Five labeling sites were chosen in this region for ^19^F NMR experiments, including Y63C, L68C and Y71C in the finger loop, T136C in the middle loop and F244C in the C loop (**Fig. 4a**). Overall, these sites adopt a homogeneous conformation in the basal state, showing a dominant major peak with linewidths in the range of about 60 ~ 120 Hz (**Fig. 4b**). The L68C and Y71C sites show smaller linewidths, indicating higher flexibility. A common spectral change observed for all five sites is the increased conformational heterogeneity upon activation. Upon V2Rpp binding, at least two different conformational states could be discriminated for the Y63C, L68C, T136C and F244C sites, whereas the L71C site shows three discernable conformations, suggesting this site displays the highest sensitivity to local structural changes. Below we analyze the spectral changes of the L71C site in more details.

**Figure 4.**
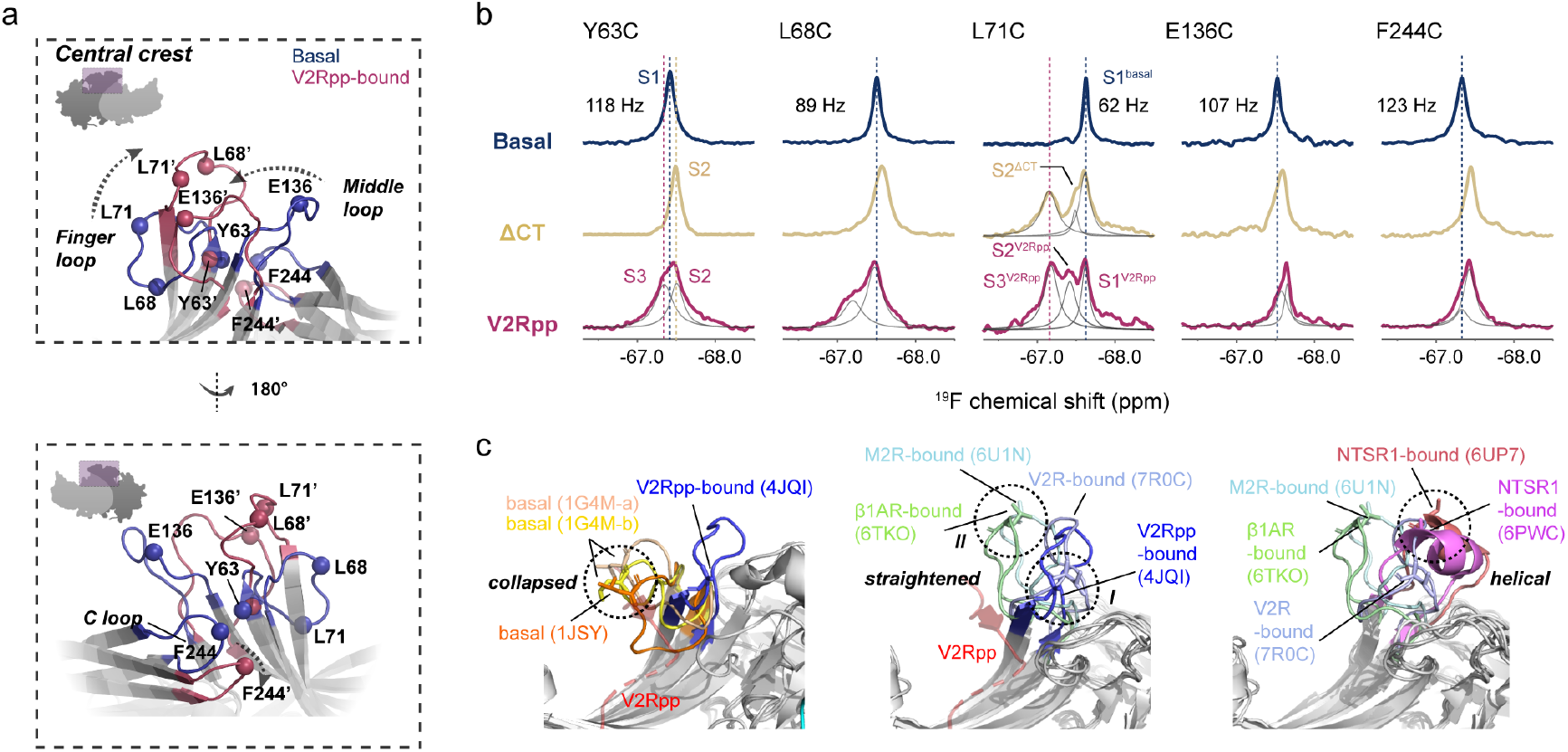
Conformational changes in the central crest probed by ^19^F NMR. (**a**) ^19^F labeling sites in the central crest mapped onto the inactive (PDB: 1JSY, blue) and V2Rpp-activated (PDB: 4JQI, magenta) βarr1 crystal structures. (**b**) ^19^F NMR spectra of residues in the central crest in the three functional states. (**c**) Different conformations of the finger loop observed in the crystal or cryo-EM structures. The L71 residue is shown in sticks in all structures. The location of the V2Rpp in the V2Rpp-bound structure (PDB: 4JQI) is depicted in red ribbon diagram in the left and middle panels.

Among the three resonances of L71C in the V2Rpp-bound state (designated as S1^V2Rpp^ to S3^V2Rpp^), S1^V2Rpp^ is essentially similar to the major peak in the basal state (S1^basal^), whereas the other two peaks show very different chemical shifts, suggesting that V2Rpp binding induces multiple conformations in the finger loop distinct from the basal state. Interestingly, L71C also shows three peaks in the ΔCT state, two of which overlay well with the S1^V2Rpp^ and S3^V2Rpp^ peaks, while the remaining one (S2^ΔCT^) shows −0.07 ppm difference in chemical shift from the S2^V2Rpp^ resonance. Therefore, CT removal is sufficient to induce finger loop activation, hallmarked by the appearance of multiple conformations, among which only the S2^ΔCT^ conformation can be further affected by V2Rpp binding.

To better interpret the multiple peaks observed in the ^19^F NMR spectra, we examined all available structures of βarr1 in the basal ^18,19^, V2Rpp-bound ^20^ and receptor-bound ^12–16^ states. The conformations of the finger loop can be grouped into three distinct categories, herein referred to as the “collapsed”, the “straightened” and the “helical” conformations (**Fig. 4c**). The “collapsed” conformation is observed in all the inactive-state structures of βarr1, in which the finger loop folds downward towards the N-domain stabilized by charge interactions between E66 in the finger loop and K292 (or K294) in the gate loop (**Supplemental data Fig. S5a**), and is most probably represented by the S1 peak in the NMR spectra. Activation of βarr1 disrupts the interaction between the finger loop and gate loop, allowing the finger loop to flip up. The “straightened” conformation is observed in the V2Rpp-bound structure (PDB: 4JQI) as well as in three receptor-bound structures, including the V2R-bound structure (PDB: 7R0C) and the M2R- and β1AR-bound structures (PDB: 6U1N and 6TKO), the latter two were ligated with V2Rpp in the C-termini to enhance binding. Interestingly, the L71 residues occupies two distinct locations in the “straightened” conformation. In the “straightened-I” conformation as observed in both the V2Rpp- and V2R-bound structures, the L71 residue is positioned close to the middle loop, whereas in the “straightened-II” conformation, observed in the M2R- and β1AR-bound structures, the finger loop further flips up to allow L71 to insert into the cytoplasmic cavity of the receptor core (**Supplemental data Fig. S5b**). Notably, L71 in the “straightened-I” conformation is also close to the 2nd phosphorylation site of V2Rpp in the V2Rpp-bound state, and this loop conformation may be further stabilized by a charge interaction between R65 and the 1st phosphorylation site (**Supplemental data Fig. S5c**), whereas such contact is incompatible with the “straightened-II” conformation due to steric clash. Therefore, we speculate that the S2 peaks in the ^19^F NMR spectra may represent the “straightened” conformations, with S2^V2Rpp^ and S2^ΔCT^ corresponding to the V2Rpp-stabilized “straightened-I” and the V2Rpp-absent “straightened-II” conformations, respectively. The downfield shift of the S2^V2Rpp^ peak compared to S2^ΔCT^ is also consistent with the de-shielding effect induced by the presence of phosphate groups. In addition, both NTSR-complexed βarr1 structures (PDB: 6UP7 and 6PWC) show a distinct finger loop conformation, in which the loop not only flips up but also forms a small helix between residues R65 and L71 (**Supplemental data Fig. S5d**). We propose that the S3 peak, which shows the largest chemical shift deviation from the basal S1 peak, may reflect a most significant local structural change and correspond to the “helical” conformation, or alternatively an ensemble of fast-exchanging conformations with higher helix-forming propensities.

Taken together, the ^19^F NMR data demonstrate that V2Rpp can sufficiently displace βarr1 CT and induce inactive-to-active transitions in the critical polar core and central crest regions. In particular, the co-existence of multiple conformational states in the finger loop observed in the activated state is in line with available structures, and could be an important feature governing the plasticity of βarrs in binding different GPCRs.

### Phosphopeptides from β2AR tail shows minimal effects on βarr1 conformation

Apart from V2Rpp, information regarding how phosphorylated tails from other receptors, especially the class A receptors, remain scarce. We herein synthesized two phosphopeptides derived from the β2AR tail, harboring GRK2- and GRK6-dependent phosphorylation patterns previously reported ^22,23^, referred to as β2AR-GRK2pp and GRK6pp hereafter, and probed their effects on βarr1 conformational dynamics (**Fig. 5a-c**).

**Figure 5.**
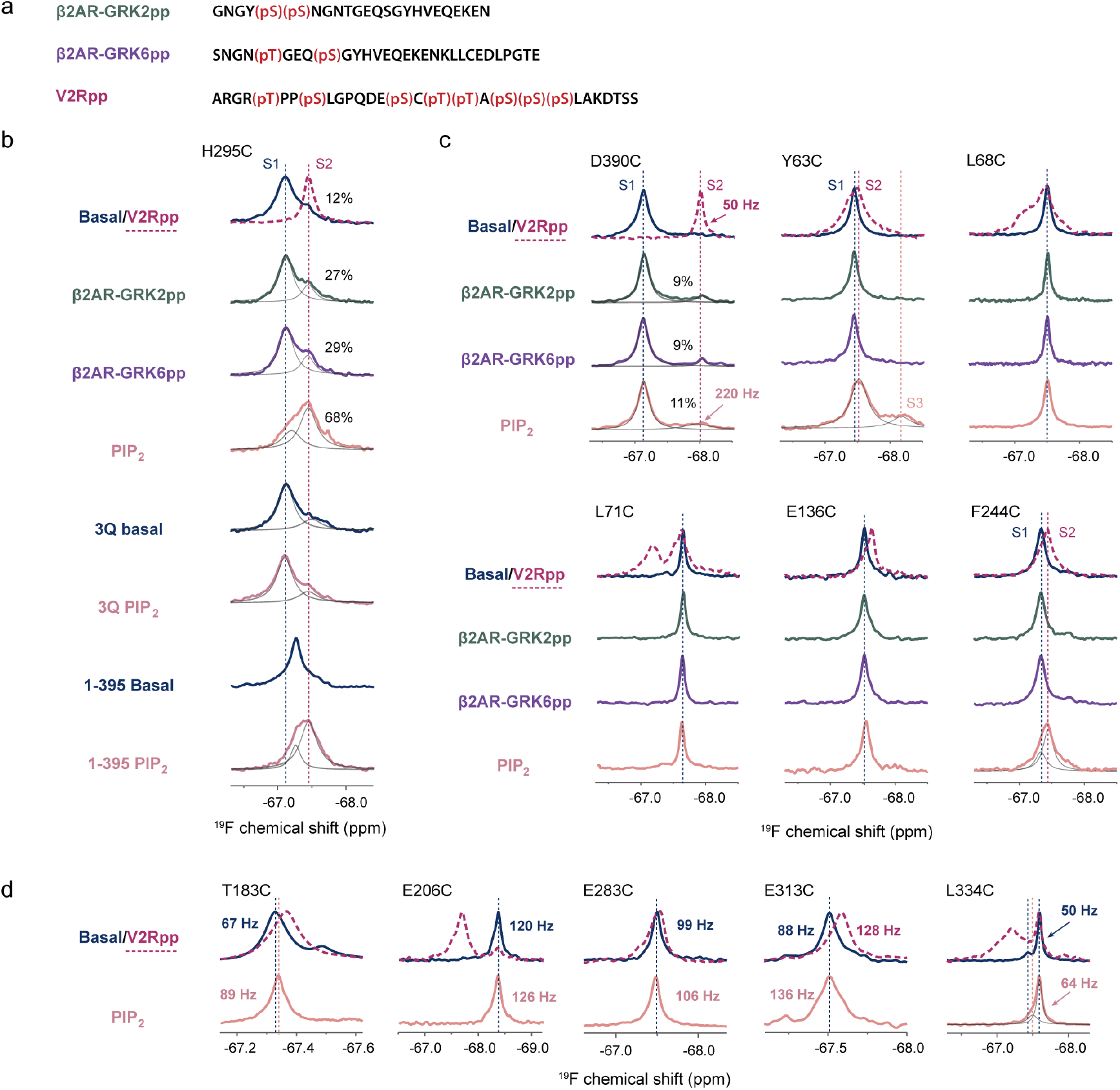
Effects of β2AR peptides and PIP_2_ binding on βarr1 conformational dynamics probed by ^19^F NMR. (**a**) Sequence and phosphorylation sites of the β2AR-GRK2pp, β2AR-GRK6pp compared with V2Rpp. (**b**) ^19^F NMR spectra of the gate loop H295C site in different states. (**c**) ^19^F NMR spectra showing the effects of β2AR-GRK2pp, -GRK6pp and PIP_2_ binding on the conformational dynamics at the CT and finger loop labeling sites. (**d**) ^19^F NMR spectra of five labeling sites in the C-domain showing the effects of PIP_2_ binding. The spectra of the V2Rpp-bound state are shown in magenta dashed lines in **c**-**d** for comparison.

Compared to V2Rpp, which fully displaces βarr1 CT at 1.2-fold molar ratio (**Fig. 2a** and **Supplemental Data Fig. S6**), the addition of 3-fold excess of either β2AR-GRK2pp or GRK6pp results in only ~ 9% fractional release of the CT as indicated by the ^19^F NMR spectra of the D390C site. Similarly, addition of excess β2AR-GRK2pp or GRK6pp induces only a small increase in the active S2 population of the H295C site, suggesting the two phosphopeptides are much less efficient in promoting conformational changes at the gate loop compared to V2Rpp. Furthermore, we observe essentially no changes in all five labeling sites in the central crest upon GRK2pp or GRK6pp-binding, indicating that these two peptides fail to induce local conformational changes associated with finger loop activation. Taken together, no obvious changes were observed in all critical structural regions upon βarr1 binding with β2AR peptides, suggesting an overall low activation level. While these observations are consistent with the functional observations that β2AR interacts transiently with βarrs in cells ^11,24–26^, question arises whether additional factors may be required for βarr1 activation.

### PIP_2_ binding modulates βarr1 conformation dynamics

Based on increasing evidence for PIP_2_ involvement in βarr activation pathway, particularly the recent report that PIP_2_-binding is required for class A receptors in βarr recruitment ^9,10^, we monitored the effect of PIP_2_ on βarr1 conformational dynamics by using its soluble derivative, diC8-PI(4,5)P2 (hereafter referred to as PIP_2_). Intriguingly, PIP_2_ binding induces broader and more significant structural responses in βarr1 than the β2AR phosphopeptides (**Fig. 5b-c**).

The most prominent change is observed in the gate loop (**Fig. 5b**). Addition of PIP_2_ alone at 1.5-fold molar ratio causes a dramatic, albeit incomplete transition to the S2 state for the H295C site, suggesting PIP_2_ is more potent in eliciting polar core activation than the β2AR tail. This partial activation effect is induced by specific PIP_2_ binding in the C-domain, because such spectral changes were not observed using a PIP_2_-binding deficient mutant βarr1-3Q (K232Q/K250Q/R236Q) ^27^. However, the very slight increase of the active S2 peak at the D390C site suggests that although PIP_2_ drastically perturbs the gate loop conformation, it is much less efficient in triggering CT release (**Fig. 5c**). Moreover, the S2 resonance in the PIP_2_-bound state shows a significantly broader linewidth (> 200 Hz) than that of the V2Rpp-bound state (~ 50 Hz), which may suggest that the PIP_2_-induced “released” conformation at the D390C site is heterogeneous or undergoes on-off exchange.

In the central crest region, PIP_2_ binding apparently perturbs Y63C and F244C while having little effect on the other sites. At the Y63C site, PIP_2_ binding not only induces the peak to shift towards the pre-activated S2 state accompanied by line broadening, as similarly observed upon V2Rpp binding, but also results in the appearance of a new minor peak S3 with large chemical shift differences from the S1 and S2 resonances, suggesting a distinct local conformation. The spectra of the F244C site, however, are similar in the PIP_2_- and V2Rpp-bound states, which can be deconvoluted into an inactive S1 and active S2 resonances, suggesting that the C-loop conformation is similarly affected by V2Rpp- or PIP_2_ binding. It is interesting to note that the Y63C and F244C sites are located at the N-, C-domain interface facing each other in the inactive state (**Supplemental Data Fig. S7**), and their concurrent spectral changes induced by PIP_2_ binding may reflect local structural rearrangement associated with interdomain rotation. Furthermore, while V2Rpp-binding most significantly perturbs the L68C and L71C sites, both positioned at the finger loop tip that flips up during activation, PIP_2_-binding perturbs only the Y63C and F244C sites located at the back side of the central crest. This apparent difference between V2Rpp- and PIP_2_-binding is consistent with the fact that they bind in the N- and C-domains, respectively, and also suggests that βarr1 can be activated through distinct mechanisms.

### Mechanism of PIP_2_-induced βarr1 activation

Our observations that PIP_2_ significantly perturbs the polar core but has limited effect on CT displacement are in agreement with previous studies ^10,28^. In particular, Janetzko *et al.* showed that PIP_2_ binding increases the active βarr1 population that can be selectively recognized by Fab30 ^10,20^. Question further arises: How does PIP_2_ binding in the C-domain activate the polar core, which is approximately 30 Å apart, without releasing the CT?

We herein examine two possible pathways by which PIP_2_-binding may be transferred to the gate loop: (1) Because the distal CT region is flexible and unobserved in the crystal structures, the possibility remains that it may transiently contact the C-domain and become perturbed by PIP_2_-binding, which then propagates through the CT towards the polar core; (2) PIP_2_-binding may induce conformational rearrangements in the C-domain structure core, and propagates through the β-strands towards the N-, C-domain interface and disturbs the polar core (**Fig. 6a**).

**Figure 6.**
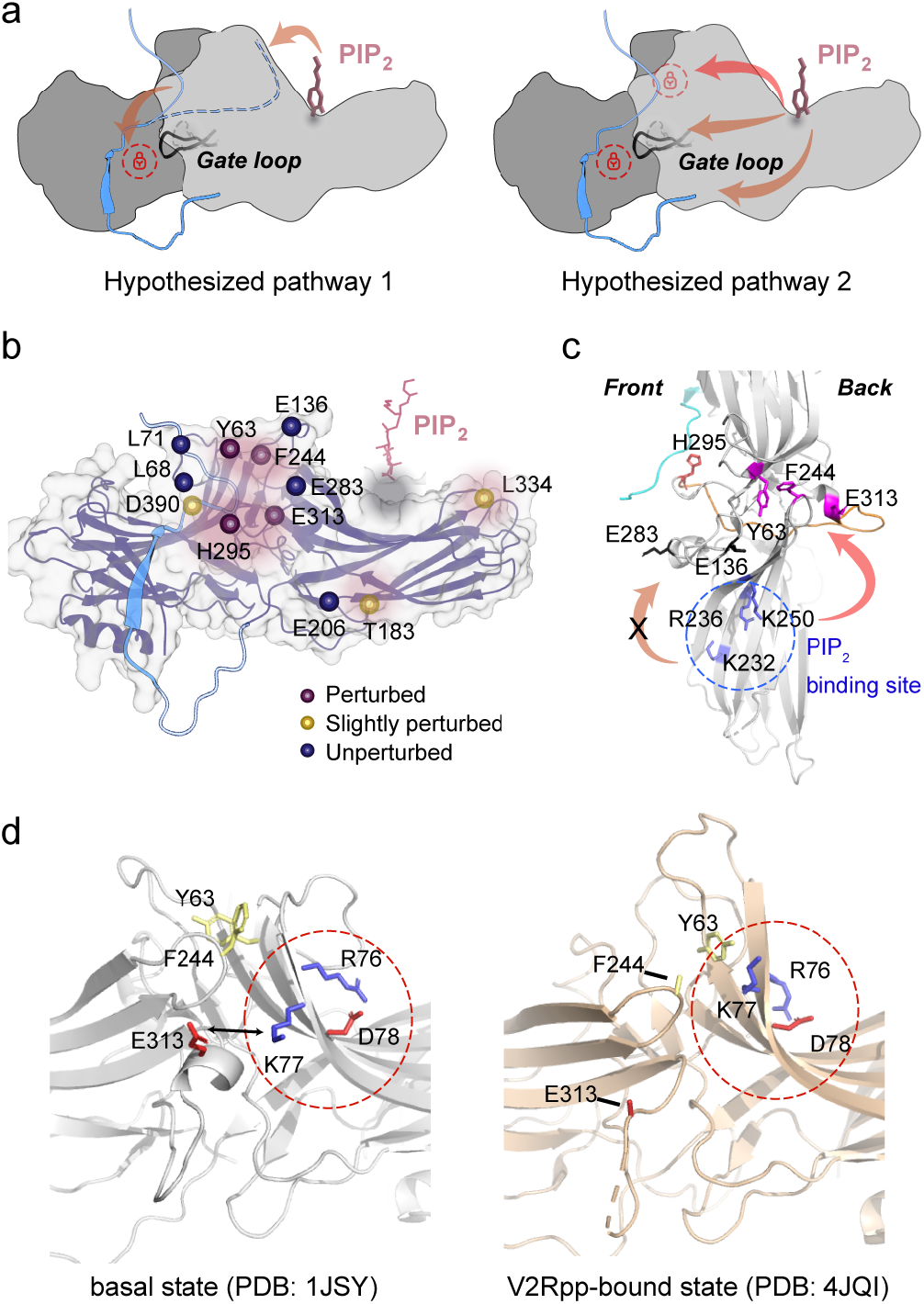
Proposed mechanism of PIP_2_-modulated βarr1 activation. (**a**) Two hypothetical pathways for PIP_2_ binding in the C-domain to be allosterically transduced to the gate loop. The most probable pathway (from the PIP_2_-binding site to the upper back region of the C-domain) is shown in a red arrow. (**b**) Summary of the effect of PIP_2_ binding on different labeling sites mapped onto the βarr1 structure. (**c**) Top view of the βarr1 structure showing the possible mechanism of PIP_2_-induced activation. The PIP_2_ binding residues are colored in blue. Residue H295 in the gate loop is colored in red. The Y63, F244 and E313 residues that are perturbed by PIP_2_ binding are colored in magenta. The E136 and E283 residues not perturbed by PIP_2_ binding are colored in black. The CT is colored in cyan. The extended loop connecting H295 and E313 is colored in orange. (**d**) Local structural comparison of the back side of the finger loop in the basal and V2Rpp-bound states. The charged residues in the finger loop proximal region are circled by red dashed lines.

We reasoned that if pathway 1 is active, deletion of the distal CT region would eliminate the effect of PIP_2_ binding on the gate loop. The H295C site of the βarr1-1-395 construct, in which the distal CT region is removed, shows a single peak in-between the S1 and S2 resonances in the basal state (**Fig. 5b**), which suggests a faster exchange between the two conformations and implies that the distal CT regions does modulate the gate loop dynamics. Nevertheless, addition of PIP_2_ can still promote the S2 conformational state to an extent comparable with full-length βarr1, indicating that pathway 1 is not responsible for PIP_2_-induced polar core activation.

To explore the 2nd pathway, we chose a few more labeling sites in the C-domain, including T183C, E206C, E283C, E313C and L334C, and examined whether they can be perturbed by PIP_2_ (**Fig. 5d & Fig. 6b-c**). The results show that PIP_2_ binding slightly perturbs both the L334C site in the C-edge loop and the T183C site in the bottom region of the C-domain, inducing ~ 30% increase of linewidths accompanied by minor peak shifts. However, very limited spectral changes are observed for the E206C or E283C sites (linewidths increase < 10%), both located closer to the N-, C-domain interface with E206 in the bottom region and E283 in the s17-s18 loop adjacent to the gate loop. In particular, E206C is dramatically affected by V2Rpp but not PIP_2_. Therefore, the PIP_2_-induced structural changes in the gate loop are not likely to be transduced through either the bottom region or the s17-s18 loop. The E313C site in the back loop, however, shows ~ 50% increase of linewidth when bound to PIP_2_, indicating significantly enhanced local dynamics. Note that E313 is located not only close to the F244 and Y63 residues, but also adjacent to a previously identified “finger-loop proximal” region that comprises a cluster of charged residues (R76, K77, D78 in βarr1 and R77, K78, D79 in βarr2) and plays a key role in locking the N-, C-domains into the inactive form ^9^. This region undergoes a substantial local structural rearrangement involving the movement of E313 away from K77 and the movement of F244 towards it (**Fig. 6d**). Moreover, previous molecular dynamics (MD) simulation suggested that K77 (K78 in βarr2) could form an interdomain salt bridge with E313 (E314 in βarr2) in the basal state, and that single mutation of either residue could result in a constitutively active phenotype of βarr2 in its accumulation in clathrin-coated endocytic structures (CCSs) ^9^. Taken together, we suggest that the NMR line-broadening of E313C upon PIP_2_ binding probably reflect a loosening of the local interdomain contact and sampling of heterogeneous conformations. This increased local dynamics unlocks the interdomain constraints from the back side of βarr1 and facilitates interdomain twisting, which is a key feature for βarr activation. The structural and dynamics changes may be transduced to the polar core through the long loop connecting H295 and E313 that extends through the βarr1 structure from front to back (**Fig. 6c**).

## DISCUSSION

Our results reveal highly complex changes of the conformational energy landscape of βarr1 during activation, which is in agreement with the previous MD study that suggested relatively independent and loosely coupled structural changes for different regions ^29^. In particular, our data demonstrate the intrinsic ability of the finger loop to adopt multiple conformations once the autoinhibitory CT is displaced, which may underlie the plasticity of βarrs in binding to diverse GPCRs. Furthermore, we show that PIP_2_ induces activation of βarr1 much more efficiently than the phosphopeptides derived from the class A receptor β2AR, and its mechanism involves local structure destabilization at the back side of the finger loop region without displacing the CT, which is consistent with previous findings ^10,28^.

Under this framework, βarr activation by GPCRs can be a more complex process involving multiple contributing factors (**Supplemental Data Fig. S8**). In the case of class B receptors, the phosphorylated tails bind to βarrs strong enough to initiate activation and form the “tail-engaged” complexes, leading to subsequent core engagement or core-independent signaling. In the case of the class A receptors, however, it remains enigmatic how such “tail-engaged” complexes are formed, and it appears that membrane lipids play a more critical role in the initial recruitment and activation of βarrs. Tail phosphorylation for class A receptors can be important since it removes the autoinhibitory effect and exposes the receptor cytoplasmic side for the core-engaged interaction as recently suggested ^28^. Based on our current results and previous studies, PIP_2_, receptor core and phosphorylated receptor tail each binds at a distinct region of βarrs, differentially destabilizing local conformations and allosterically affecting more distal areas. These binding events can act as independent signal inputs, and their combinatory effects help reshape the conformational energy landscape of βarrs, leading to full activation as well as diverse functional outcomes. Given the diverse receptor tail phosphorylation patterns and the presence of other types membrane lipids, we anticipate the regulation mechanism of the βarr signaling pathway to be extremely multifaceted and intricate.

## METHODS

### Constructs

The cysteine-less mutant (C59V/C125S/C140L/C150V/C242V/C251V/C269S) of the long splice variant of human βarr1 was used for introduction of single site cysteine mutation for ^19^F labeling. The sequence was codon-optimized for expression in *Escherichia coli* and cloned into a pET-28a(+) vector (Novagen), with an N-terminal 6xHis-tag, followed by an SSG linker and a thrombin cleavage site. All mutants, including the 1-382 (βarr1-ΔCT) and 1-395 truncation constructs, were generated using site-directed mutagenesis kits (Thermo Fisher).

### Expression and purification of βarr1

All constructs of βarr1 were transformed into *E. coli* BL21(DE3) cells. Cells were cultured in 20 mL Luria Bertani (LB) medium containing 50 mg/L kanamycin at 37 °C for 12 h and then transferred into Terrific Broth (TB) medium for large-scale cultivation at 37 °C. When OD600 reached 1.0, the cells were transferred to 18 °C and shaked for 1h before induction with 50 μM isopropyl-β-D-thiogalactopyranoside (IPTG). After 24h of induction, cells were harvested, centrifuged, resuspended in the lysis buffer (50 mM Tris-HCl, 500 mM NaCl, 15% glycerol, pH 8.0) to a final volume of 40 ml per liter of cells, and then flash-frozen with liquid nitrogen and stored at −80 °C.

During purification of βarr1, all procedures describe below were performed at 4 °C or on ice. Cells were lysed by sonication, supplemented with protease inhibitor cocktail. The supernatant of the lysis was loaded onto a Ni-IDA Sepharose column equilibrated with the lysis buffer, and subsequently washed using buffer A (20 mM Tris-HCl, 500 mM NaCl, 10% glycerol, 40 mM imidazole, pH 8.0) and buffer B (20 mM Tris-HCl, 150 mM NaCl, 40 mM imidazole, pH 8.0). The target protein was eluted with 10 column volumes of buffer C (20 mM Tris-HCl, 500 mM NaCl, 5% glycerol, 250 mM imidazole, pH 8.0). The eluted protein was concentrated to 10 mg/ml and exchanged into buffer D (20 mM Tris-HCl, 500 mM NaCl, 5% glycerol, pH 8.0) by repeated centrifugation and dilution using a Millipore concentrator with 30-kDa molecular weight cut-off. Thrombin was added into the protein sample at 1:50 w/w concentration and incubated overnight to ensure complete cleavage of the His-tag, which was verified by SDS-PAGE. Then the protein was applied again to the Ni-IDA column and the elute was collected and exchanged to buffer E (20 mM Tris-HCl, 150 mM NaCl, pH 8.0). The protein was finally purified by size-exclusion chromatography using a Superdex 200 Increase 10/300 GL column (GE Healthcare) with buffer E and concentrated to 10 mg/ml, supplemented with leupeptin and bestatin (Sigma-Aldrich) to inhibit protease activity, flash-frozen with liquid nitrogen and stored at −80 °C until use. The purities of all protein samples were verified by SDS-PAGE. One-dimensional ^1^H NMR and circular dichroism spectroscopy were employed to verity that all samples were correctly folded.

### ^19^F labeling of βarr1

Each single-cysteine mutant of βarr1 was diluted to a concentration of 50 μM to be labeled using the sulfoxide-based ^19^F probe wPSP-6F, which was derived from the previously reported probes ^30^. The wPSP-6F stock dissolved in buffer E was added into the protein sample at 1.5-fold molar ratio excess and incubated at 4 °C for 1 h, followed by quenching with 2-fold molar ratio of dithiothreitol. The sample was then fully exchanged using buffer E to remove excess of small molecules, and subjected to a second round of SEC purification. The purified proteins were concentrated to 100 μM, flash-frozen in liquid nitrogen and stored at −80 °C until use. The purities of all samples were verified by SDS-PAGE.

### Peptides

All phosphopeptides used in this study were obtained from custom peptide synthesis (Scilight-peptide). The purities of the peptides were verified by analytical high-performance liquid chromatography with > 95% purity. The peptides were weighted and dissolved in buffer E at concentrations of 1 mM (V2Rpp) or 3 mM (β2AR-GRK2pp and -GRK6pp). The exact concentrations of the V2Rpp and β2AR-GRK6pp peptide stocks were determined by cysteine reaction with the Ellman’s reagent as previously described ^29^, cross-validated with UV absorbance at 205 nm ^31^ or 280 nm. The exact concentration of the β2AR-GRK2pp, which contains no cysteine residue, was determined by UV absorbance at 280 nm and cross-validated by absorbance at 205 nm.

### Preparation of βarr1 samples in different states

The ^19^F-labeled βarr1 was diluted to 50 μM using buffer E before incubation with peptides or PIP_2_. The V2Rpp, β2AR-GRK2pp/GRK6pp or PIP_2_ were added at 1.2-fold, 3-fold and 1.5-fold molar ratio excess to the protein, respectively. All complexes were incubated at 25 °C for 1 h, and the freshly prepared samples were immediately subjected to ^19^F NMR experiments.

### NMR experiments

NMR samples were prepared in buffer E to a final volume of 140 μL, supplemented with 1 mM DTT, 10% D_2_O for field lock, and 1 μM sodium trifluoroacetate (STFA) as the internal chemical shift reference. All samples were sterile-filtered and loaded into sterile 3 mm NMR microtubes to prevent microbial contamination. Final protein concentrations vary for different constructs, depending on their expression levels and stabilities, but all in the range of 20-50 μM.

All spectra were recorded at 25 °C using a Bruker AVANCE III HD 800 MHz spectrometer equipped with a ^1^H-^19^F/^13^C/^15^N TCI triple resonance cryogenic probe (Bruker). For all ^19^F NMR experiments, the carrier frequency was set to −68.0 ppm and the spectra were recorded with 4k points and a spectral width of 15000 Hz. For signal optimization, the experiments were conducted with a 0.8 ms pre-scan-delay and a 100 ms relaxation delay, and acquired using 100000-200000 scans, yielding a signal-to-noise (S/N) ratio of approximately 50. SDS-PAGE were run for all samples before and after NMR experiments to ensure that no protein degradation occurs during the experiments.

### NMR data analysis

All NMR data were analyzed using the MestRaNova 12.0.0 software (https://mnova.pl). The spectra were baseline-corrected by the Whittaker Smoother method and applied with a 30 Hz exponential windows function. The ^19^F chemical shifts were referenced to STFA at −75.450 ppm. All datasets were processed using the same variables to facilitate comparison. Spectral deconvolutions were performed assuming generalized Lorentzian line shapes. To distinguish overlapping resonances, a line broadening (LB) value of 3 Hz was initially applied to ensure the highest resolution and determine the exact chemical shift values. The spectra were subsequently fitted iteratively using an LB value of 30 Hz. Peak intensities, integrated peak area and linewidths were determined based on the fitted results.

### Clathrin binding assay

Binding of βarr1 to clathrin was monitored by glutathione S-transferase (GST) pull-down assay using the GST-fused clathrin terminal domain (1-363) as a bait, similar to previously described ^32,33^. Briefly, human clathrin terminal domain (clatrin-TD) was cloned into a pET-21a(+) vector (Novagen), with an N-terminal GST-tag. The GST-clatrin-TD was expressed in *E. coli* BL21(DE3) cells and purified by glutathione-sepharose 4B beads (GE Healthcare) following previously reported protocols ^33^. For the GST pull-down experiments, 4 μM wild-type or mutant βarr1 was mixed with or without 4 μM V2Rpp in 40 μL pull-down buffer (50 mM Tris, 150 mM NaCl, 2 mM EDTA, 5 mM DTT, pH 8.0), and incubated at 4 °C for 30 min. Stock solution of GST-clathrin was subsequently added into the mixtures to a final concentration of 2 μM and incubated for another 30 min at 4 °C. Then, the samples were supplemented with 15 μL glutathione agarose beads, diluted to a final volume of 200 μL and incubated at 4 °C for 5 h. The beads were collected by centrifugation and washed using the pull-down buffer. After removing the supernatant, the beads were mixed with 15 μL 2× SDS loading buffer and boiled for 10 min. Proteins were separated by SDS-PAGE gel and visualized by Coomassie blue staining.

## ACKNOWLEDGEMENTS

All NMR experiments were performed at the Beijing NMR Center and the NMR facility of National Center for Protein Sciences at Peking University. This work was supported by funding from the National Key R&D Program of China (2016YFA0501201) and the National Natural Science Foundation of China (21991083). Y. H. acknowledges financial supports from the National Center for Protein Sciences at Peking University (KF-202005) and the Huanghe Talents Plan from Wuhan city.

## AUTHOR CONTRIBUTIONS

R.Z., C.J. and Y.H. designed the research. R.Z. prepared the samples, performed the NMR experiments and analyzed the data. Z.W., Z.C. and C.L. provided ^19^F-labeling probes. R.Z., C.J. and Y.H. provided structural interpretations of the experimental data. R.Z. and Y.H. prepared the figures. Y.H. wrote the paper with input from all authors.

## REFERENCES

1 Luttrell, L. M. & Lefkowitz, R. J. The role of beta-arrestins in the termination and transduction of G-protein-coupled receptor signals. J Cell Sci 115, 455–465, doi:10.1242/jcs.115.3.455 (2002).

2 Caron, M. G. & Barak, L. S. A Brief History of the β-Arrestins. Methods Mol Biol 1957, 3–8, doi:10.1007/978-1-4939-9158-7_1 (2019).

3 Lefkowitz, R. J. & Shenoy, S. K. Transduction of receptor signals by beta-arrestins. Science 308, 512–517, doi:10.1126/science.1109237 (2005).

4 DeWire, S. M., Ahn, S., Lefkowitz, R. J. & Shenoy, S. K. Beta-arrestins and cell signaling. Annu Rev Physiol 69, 483–510, doi:10.1146/annurev.physiol.69.022405.154749 (2007).

5 Shukla, A. K. et al. Visualization of arrestin recruitment by a G-protein-coupled receptor. Nature 512, 218–222, doi:10.1038/nature13430 (2014).

6 Thomsen, A. R. B. et al. GPCR-G Protein-β-Arrestin Super-Complex Mediates Sustained G Protein Signaling. Cell 166, 907–919, doi:10.1016/j.cell.2016.07.004 (2016).

7 Jala, V. R., Shao, W. H. & Haribabu, B. Phosphorylation-independent beta-arrestin translocation and internalization of leukotriene B4 receptors. J Biol Chem 280, 4880–4887, doi:10.1074/jbc.M409821200 (2005).

8 Jung, S. R., Kushmerick, C., Seo, J. B., Koh, D. S. & Hille, B. Muscarinic receptor regulates extracellular signal regulated kinase by two modes of arrestin binding. Proc Natl Acad Sci U S A 114, e5579–e5588, doi:10.1073/pnas.1700331114 (2017).

9 Eichel, K. et al. Catalytic activation of β-arrestin by GPCRs. Nature 557, 381–386, doi:10.1038/s41586-018-0079-1 (2018).

10 Janetzko, J. et al. Membrane phosphoinositides regulate GPCR-β-arrestin complex assembly and dynamics. Cell 185, 4560–4573.e4519, doi:10.1016/j.cell.2022.10.018 (2022).

11 Oakley, R. H., Laporte, S. A., Holt, J. A., Barak, L. S. & Caron, M. G. Molecular determinants underlying the formation of stable intracellular G protein-coupled receptor-beta-arrestin complexes after receptor endocytosis*. J Biol Chem 276, 19452–19460, doi:10.1074/jbc.M101450200 (2001).

12 Bous, J. et al. Structure of the vasopressin hormone-V2 receptor-β-arrestin1 ternary complex. Sci Adv 8, eabo7761, doi:10.1126/sciadv.abo7761 (2022).

13 Yin, W. et al. A complex structure of arrestin-2 bound to a G protein-coupled receptor. Cell Res 29, 971–983, doi:10.1038/s41422-019-0256-2 (2019).

14 Huang, W. et al. Structure of the neurotensin receptor 1 in complex with β-arrestin 1. Nature 579, 303–308, doi:10.1038/s41586-020-1953-1 (2020).

15 Lee, Y. et al. Molecular basis of beta-arrestin coupling to formoterol-bound beta1-adrenoceptor. Nature 583, 862–866, doi:10.1038/s41586-020-2419-1 (2020).

16 Staus, D. P. et al. Structure of the M2 muscarinic receptor-β-arrestin complex in a lipid nanodisc. Nature 579, 297–302, doi:10.1038/s41586-020-1954-0 (2020).

17 Hanson, S. M. et al. Arrestin mobilizes signaling proteins to the cytoskeleton and redirects their activity. J Mol Biol 368, 375–387, doi:10.1016/j.jmb.2007.02.053 (2007).

18 Han, M., Gurevich, V. V., Vishnivetskiy, S. A., Sigler, P. B. & Schubert, C. Crystal structure of beta-arrestin at 1.9 A: possible mechanism of receptor binding and membrane Translocation. Structure 9, 869–880, doi:10.1016/s0969-2126(01)00644-x (2001).

19 Milano, S. K., Pace, H. C., Kim, Y. M., Brenner, C. & Benovic, J. L. Scaffolding functions of arrestin-2 revealed by crystal structure and mutagenesis. Biochemistry 41, 3321–3328, doi:10.1021/bi015905j (2002).

20 Shukla, A. K. et al. Structure of active β-arrestin-1 bound to a G-protein-coupled receptor phosphopeptide. Nature 497, 137–141, doi:10.1038/nature12120 (2013).

21 Kim, Y. J. et al. Crystal structure of pre-activated arrestin p44. Nature 497, 142–146, doi:10.1038/nature12133 (2013).

22 Yang, F. et al. Phospho-selective mechanisms of arrestin conformations and functions revealed by unnatural amino acid incorporation and (19)F-NMR. Nat Commun 6, 8202, doi:10.1038/ncomms9202 (2015).

23 Nobles, K. N. et al. Distinct phosphorylation sites on the β(2)-adrenergic receptor establish a barcode that encodes differential functions of β-arrestin. Sci Signal 4, ra51, doi:10.1126/scisignal.2001707 (2011).

24 Oakley, R. H., Laporte, S. A., Holt, J. A., Caron, M. G. & Barak, L. S. Differential affinities of visual arrestin, beta arrestin1, and beta arrestin2 for G protein-coupled receptors delineate two major classes of receptors. J Biol Chem 275, 17201–17210, doi:10.1074/jbc.M910348199 (2000).

25 Zhang, J. et al. Cellular trafficking of G protein-coupled receptor/beta-arrestin endocytic complexes. J Biol Chem 274, 10999–11006, doi:10.1074/jbc.274.16.10999 (1999).

26 Oakley, R. H., Laporte, S. A., Holt, J. A., Barak, L. S. & Caron, M. G. Association of beta-arrestin with G protein-coupled receptors during clathrin-mediated endocytosis dictates the profile of receptor resensitization. J Biol Chem 274, 32248–32257, doi: 10.1074/jbc.274.45.32248 (1999).

27 Gaidarov, I., Krupnick, J. G., Falck, J. R., Benovic, J. L. & Keen, J. H. Arrestin function in G protein-coupled receptor endocytosis requires phosphoinositide binding. EMBO J 18, 871–881, doi:10.1093/emboj/18.4.871 (1999).

28 Asher, W. B. et al. GPCR-mediated β-arrestin activation deconvoluted with single-molecule precision. Cell 185, 1661–1675.e1616, doi:10.1016/j.cell.2022.03.042 (2022).

29 Latorraca, N. R. et al. How GPCR Phosphorylation Patterns Orchestrate Arrestin-Mediated Signaling. Cell 183, 1813–1825.e1818, doi:10.1016/j.cell.2020.11.014 (2020).

30 Chai, Z. et al. Simultaneous detection of small molecule thiols with a simple (19)F NMR platform. Chem Sci 12, 1095–1100, doi:10.1039/d0sc04664g (2020).

31 Anthis, N. J. & Clore, G. M. Sequence-specific determination of protein and peptide concentrations by absorbance at 205 nm. Protein Sci 22, 851–858, doi:10.1002/pro.2253 (2013).

32 Kang, D. S. et al. Structure of an arrestin2-clathrin complex reveals a novel clathrin binding domain that modulates receptor trafficking. J Biol Chem 284, 29860–29872, doi:10.1074/jbc.M109.023366 (2009).

33 Kim, Y. M. & Benovic, J. L. Differential roles of arrestin-2 interaction with clathrin and adaptor protein 2 in G protein-coupled receptor trafficking. J Biol Chem 277, 30760–30768, doi:10.1074/jbc.M204528200 (2002).

