## Supplemental data for "Distinct activation mechanisms of β-arrestin 1 revealed by ^19^F NMR spectroscopy"

### Supplemental Data Figure S1-S8 for "Distinct activation mechanisms of β-arrestin 1 revealed by $^{19}$F NMR spectroscopy"

Ruibo Zhai, Zhuoqi Wang, Zhaofei Chai, Conggang Li, Changwen Jin, Yunfei Hu

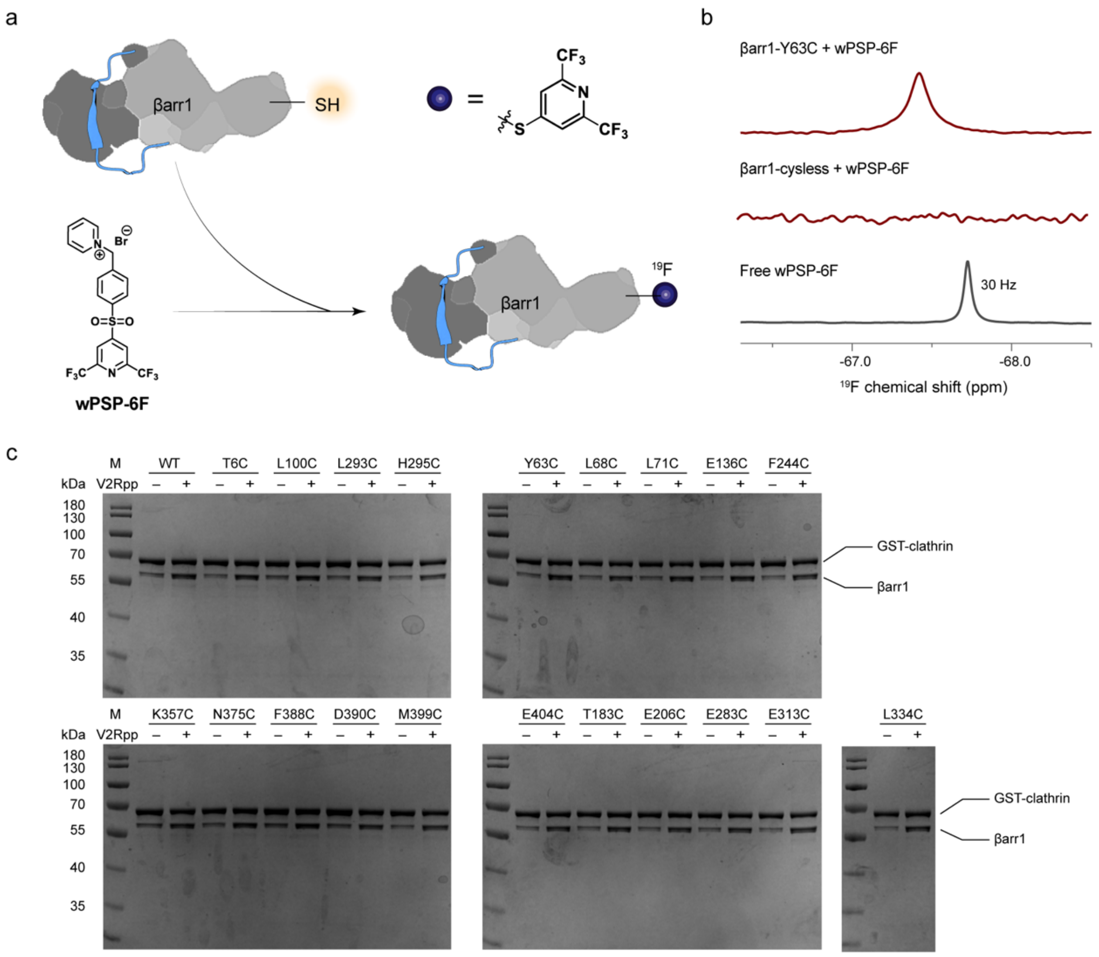


**Figure S1. $^{19}$F-labeling scheme for βarr1.** (**a**) A schematic illustration showing the single-site labeling of $^{19}$F-probe by reaction with a free cysteine thiol group. (**b**) Representative spectra of βarr1 showing the verification of successful $^{19}$F labeling, and comparison with the $^{19}$F spectrum of the free $^{19}$F probe. (**c**) GST pull-down assays showing the V2Rpp-enhanced binding to clathrin by wild-type and mutant βarr1 proteins.

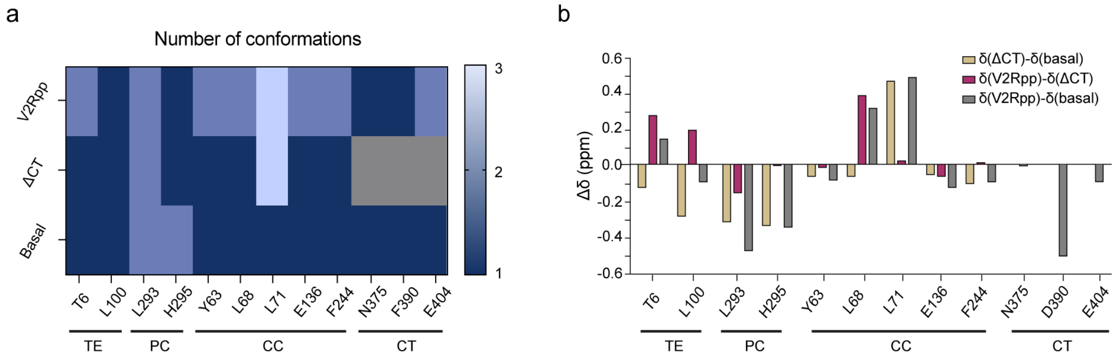


**Figure S2. V2Rpp-induced conformational changes in critical regions of βarr1.** (**a**) Summary of the number of conformational states in key structural regions in the basal, ΔCT and V2Rpp-bound states of βarr1. (**b**) Chemical shift changes for representing residues in key structural regions in the basal, ΔCT and V2Rpp-bound states of βarr1. The TE, PC, CC, CT abbreviations stand for the three-element interacting site, the polar core, the central crest and the carboxyl tail, respectively. For residues showing multiple peaks, only the largest chemical shift changes are presented in (b).

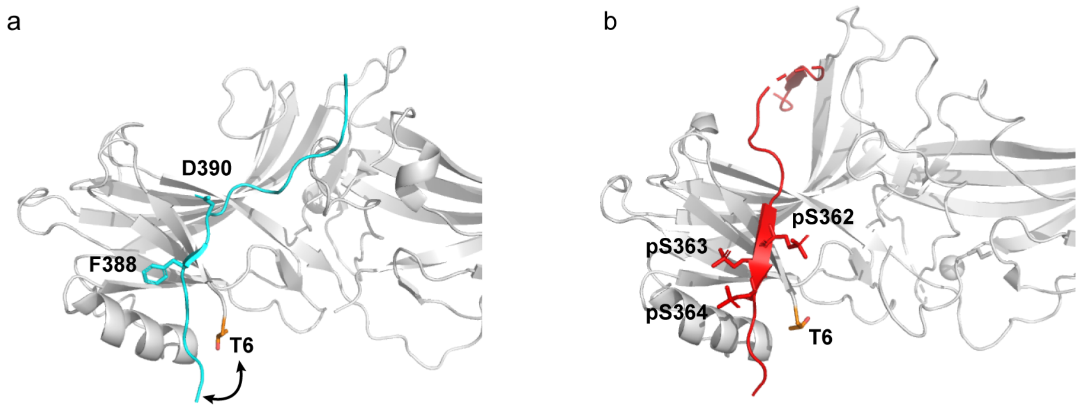


**Figure S3. Local structure comparison at the TE interacting region of βarr1.** (**a**) The location of T6 in the inactive state structure of βarr1 (PDB: 1JSY). The CT region is colored in cyan and the side chains of F388 and D390 are shown as sticks. (**b**) The location of T6 in the structure of V2Rpp-activatedβarr1 (PDB: 4JQI). The bound V2Rpp is colored in red, and the cluster of triple phosphoserine residues are shown as sticks.

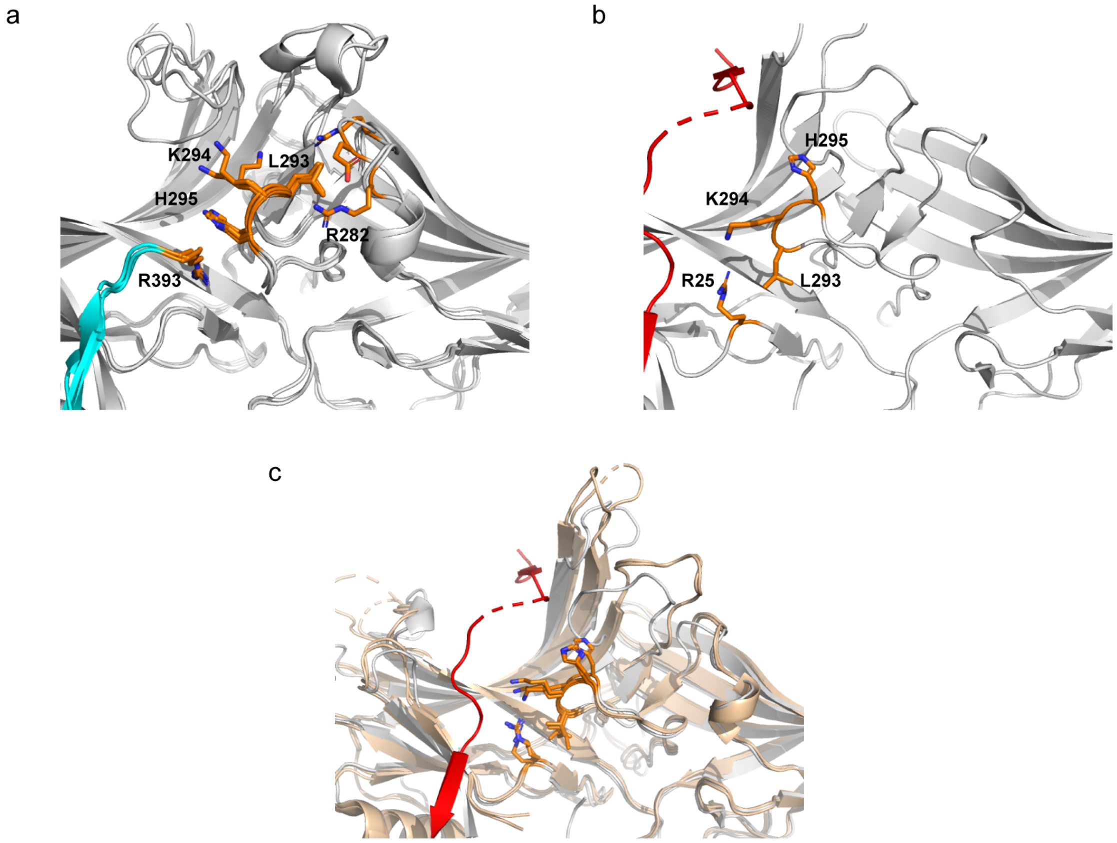


**Figure S4. Local structure of the gate loop observed in crystal structures.** (**a**) Gate loop conformations observed in the inactive state crystal structures, including PDB 1JSY and both molecules in PDB 1G4M. The CT region is shown in cyan. (**b**) Gate loop conformations observed in the V2Rpp-bound active state crystal structure (PDB: 4JQI). The V2Rpp is shown in red. (**c**) Comparison of the gate loop conformations in the V2Rpp-bound βarr1 structure (PDB: 4JQI, white) and the CT-truncated splice variant p44 of visual arrestin (both molecules in PDB 4J2Q, wheat).

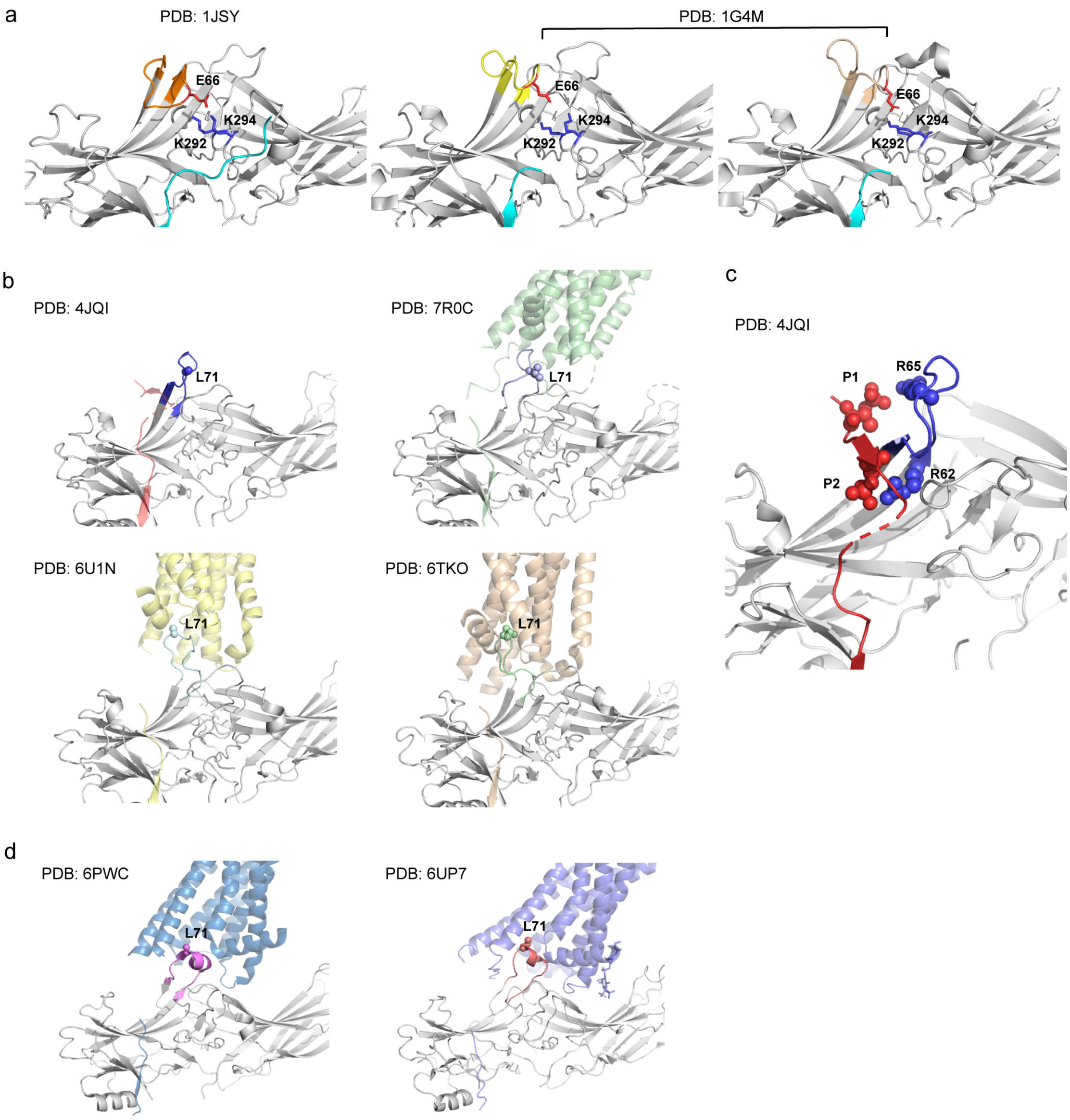


**Figure S5. Local structure of the finger loop observed in crystal/cryo-EM structures.** (**a**) The "collapsed" conformation of the finger loop observed in the inactive state crystal structures. (**b**) The two types of "extended" conformations of the finger loop observed in the V2Rpp-bound and receptor-bound structures. (**c**) Contacts between the 1st and 2nd phosphorylation site (P1 and P2) in the V2Rpp and the finger loop residues observed in the V2Rpp-bound βarr1 structure (PDB: 4JQI). (**d**) The "helical" conformation of the finger loop observed in receptor-bound structures.

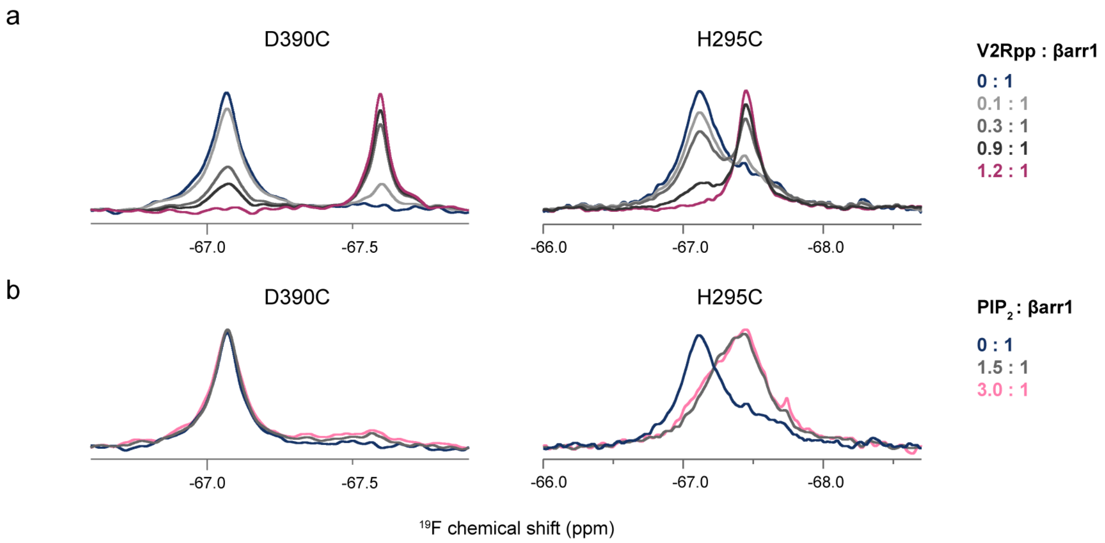


**Figure S6. Titration experiments of V2Rpp or $PIP_2$ into βarr1 probed by $^{19}F$ NMR.** (**a**) Titration of V2Rpp up to 1.2:1 molar ratio activates βarr1 at both D390C and H295C sites. (**b**) Titration of $PIP_2$ up to 3.0:1 molar ratio partially activates βarr1 at the H295C site, but minimally affects the D390C site.

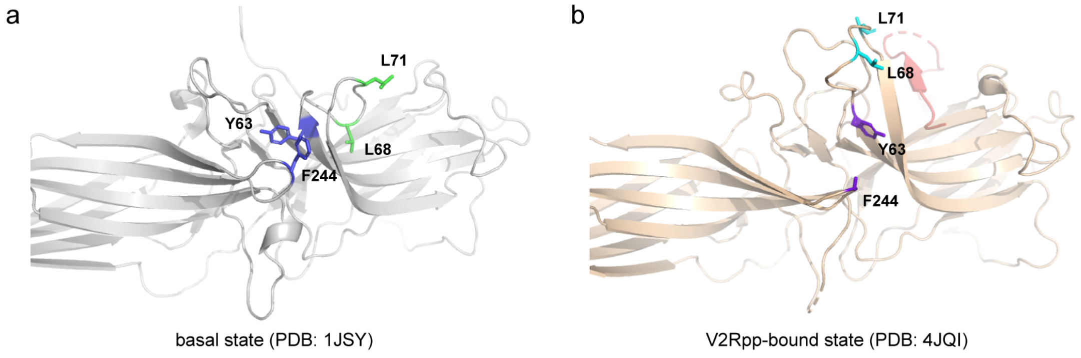


**Figure S7. Local structure showing the back site of the central crest region observed in βarr1 crystal structures.** (**a**) Local structure showing the packing between Y63 and F244 in the basal state. (**b**) Local structure showing the break of the Y63-F244 contact in the V2Rpp-activated state due to interdomain rotation. The side chain electron density of F244 is missing and therefore not displayed. The Y63 and F244 residues are colored in blue and purple, while the L68 and L71 residues are colored in green and cyan in the two states, respectively. The V2Rpp is shown in red in (**b**).

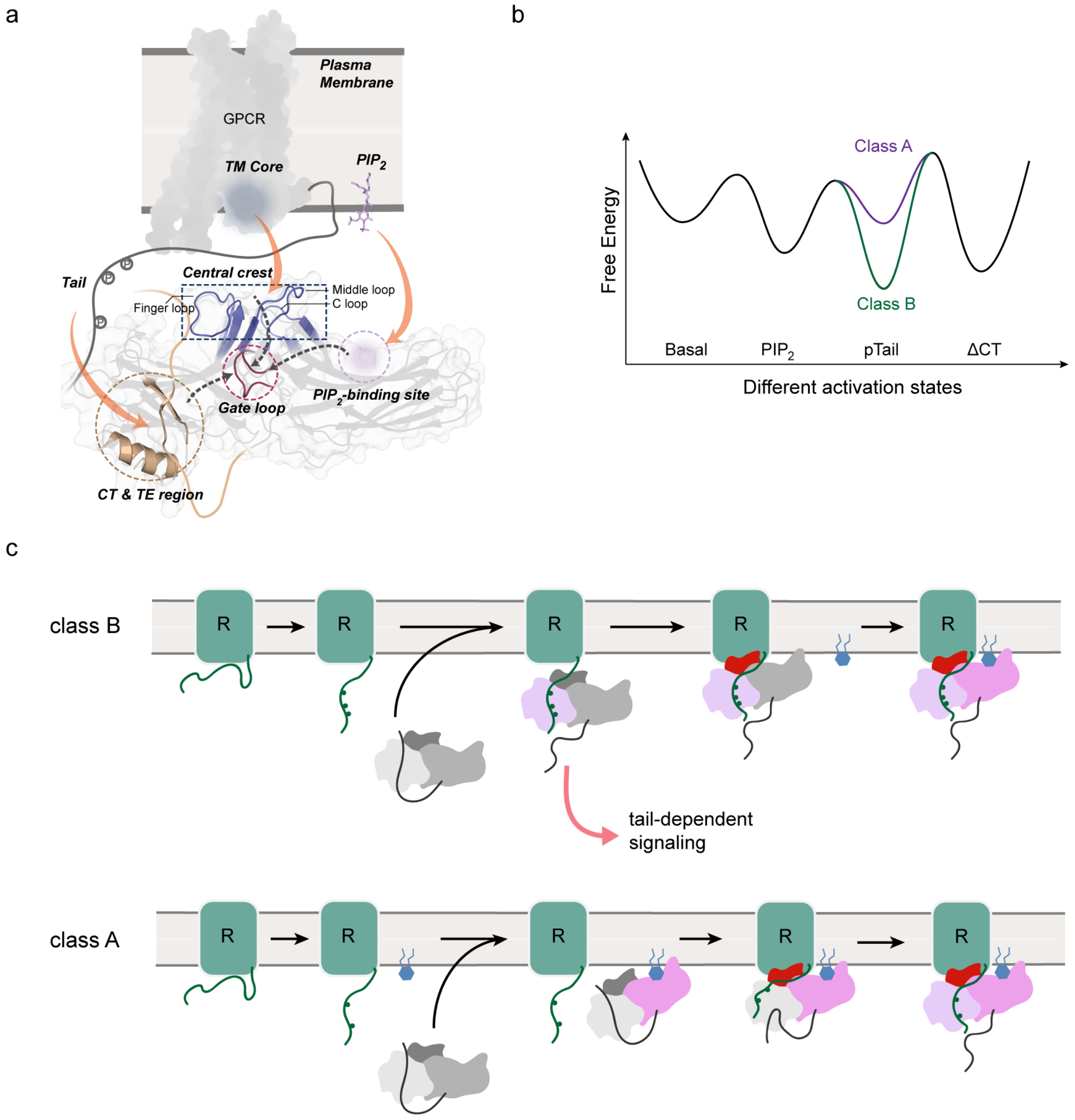


**Figure S8. Hypothesis for different activation mechanisms of βarr1.** (**a**) A schematic illustration showing the binding of receptor tail, receptor TM core and membrane $PIP_2$ to different structural regions of βarr1 (orange arrows), and transduction of conformational changes towards the gate loop, a central hub undergoing substantial rearrangements during activation. (**b**) An illustrative diagram showing the modulation of βarr1 conformational energy landscape by binding of $PIP_2$ and the phosphorylated tails (pTail) of difference receptor classes. (**c**) Schematic illustrations showing the hypothesized βarr1 activation process by class B and class A receptors. The color changes in different regions of βarr1 (the N-domain, the C-domain and the central crest) represent their individual binding and structural activation events.
